# Elucidating the Glycan-Binding Specificity and Structure of *Cucumis melo* Agglutinin, a New R-Type Lectin

**DOI:** 10.1101/2023.11.30.569503

**Authors:** Jon Lundstrøm, Emilie Gillon, Valérie Chazalet, Nicole Kerekes, Antonio Di Maio, Yan Liu, Ten Feizi, Annabelle Varrot, Daniel Bojar

## Abstract

Plant lectins have garnered attention for their roles as laboratory probes and potential therapeutics. Here, we report the discovery and characterization of *Cucumis melo* agglutinin (CMA1), a new R-type lectin from melon. Our findings reveal CMA1’s unique glycan-binding profile, mechanistically explained by its 3D structure, augmenting our understanding of R-type lectins. We expressed CMA1 recombinantly and assessed its binding specificity using multiple glycan arrays, covering 1,046 unique sequences. This resulted in a complex binding profile, strongly preferring C2-substituted, beta-linked galactose (both GalNAc and Fuca1-2Gal), which we contrasted with the established R-type RCA1 lectin. We also report binding of specific glycosaminoglycan subtypes and a general enhancement of binding by sulfation. Further validation using agglutination, thermal shift assays, and surface plasmon resonance confirmed and quantified this binding specificity in solution. Finally, we solved the high-resolution structure of the CMA1 N-terminal domain using X-ray crystallography, supporting our functional findings at the molecular level. Our study provides a comprehensive understanding of CMA1, laying the groundwork for further exploration of its biological and therapeutic potential.

## Introduction

Lectins have long been the subject of intense scientific scrutiny, serving as molecular bridges that span the realms of biochemistry, cellular biology, and biomedicine. These carbohydrate-binding proteins boast a range of functions, acting as recognition modules in cell-molecule and cell-cell interactions, thereby playing vital roles in immune defense, regulation of growth, and apoptosis^1^. In plants, they serve as essential components in development, immunity, and stress signaling^2,3^.

In light of the burgeoning interest in the intersection of glycobiology and biomedicine, the characterization of new lectins has carved out a significant niche in scientific research. Specifically, lectins have emerged as invaluable tools for staining cells and tissues, thereby offering insights into cellular heterogeneity and function. For instance, the use of wheat germ agglutinin (WGA) and concanavalin A (ConA) has been instrumental in selectively staining cells based on their glycan expression^4^, including single-cell approaches^5,6^. In the realm of therapeutics, lectins such as mistletoe lectins have shown promise in cancer therapy, by virtue of their ability to induce apoptosis in malignant cells^7^. Further, the creation of lectin arrays^8,9^, which employ a diverse set of characterized lectins, has enabled high-throughput glycan profiling, thereby advancing both diagnostic methods and biomarker discovery. Examples include arrays that can rapidly profile alterations in glycosylation patterns, pivotal in many diseases and inflammatory changes^10,11^.

Many of the most commonly used lectins for the abovementioned applications are R-type lectins, especially those derived from plants. Examples include SNA (from *Sambucus nigra*, binding Neu5Acα2-6^12^) or RCA1 (from *Ricinus communis*, binding terminal β-linked galactose^13^).

Yet, despite the extensive studies on plant lectins, particularly R-type lectins, there are still significant gaps in our understanding. Further, in general, few melon lectins have been studied in detail. Some reports indicate the presence of chitooligosaccharide-binding (i.e., β1-4 GlcNAc oligomers) lectins from phloem exudates of melons^14,15^, as well as R-type lectins in bitter melon^16^, yet not much else is known about binding specificities exhibited by lectins derived from melons. In particular, existing research in this area often lacks a comprehensive characterization that includes both functional and structural analysis of these lectins.

Here, we introduce a novel member of characterized melon lectins: the *Cucumis melo* agglutinin (CMA1), an R-type lectin derived from melon. Prior to our study, CMA1 was only a predicted protein from genomic sequencing, with moderate certainty scores on lectin-specific databases. Our comprehensive analysis using glycan array experiments, thermal shift assays, and high-resolution X-ray crystallography not only confirms its classification as a functional R-type lectin but also provides a deep dive into its unique glycan-binding profile and high-resolution 3D structure. Overall, we present a deeply characterized new lectin with a unique binding profile of specifically recognizing C2-substituted galactose.

## Results and Discussion

### The identification and production of a new lectin from the melon *Cucumis melo*

CMA1 is a predicted protein from whole-genome shotgun sequencing of leaves from the melon plant *Cucumis melo* (variant *makuwa*, taxon ID: 1194695)^17^ and has, to our knowledge, never been studied before. With prediction scores of 0.453 on LectomeXplore^18^ and 0.251 on TrefLec^19^ (from 0, lowest, to 1, highest), CMA1 is moderately certain in its prior classification as a lectin. CMA1 comprises 291 amino acids and is predicted to fold into two linked β-trefoil domains belonging to Carbohydrate-Binding Module Family 13 (CBM13) and placing it into the group of R-type lectins. Both CBM13 domains are likely to exhibit carbohydrate-binding activity due to the conservation of key amino acids in at least one of the three potential binding sites. In contrast to other R-type lectins such as ricin, it lacks a catalytic domain.

As R-type lectins are both a well-investigated family of lectins and widely used in research and beyond, we first wanted to analyze where it would be situated in the broader context of R-type lectins. A multiple sequence alignment of binding domains of representative R-type lectins (Fig. 1a) showed that CMA1 exhibited a binding domain with a relatively similar sequence as those of the plant lectins SNA and ricin. However, we note that, in general, the substantial heterogeneity of binding motifs of even closely related lectins (SNA: Neu5Acα2-6, ricin: Gal/GalNAc) does not allow for a strong a priori hypothesis of what CMA1 would bind, even though R-type lectins in general are thought to prefer the Gal/GalNAc type motif mentioned in the context of ricin^20^.

**Figure 1.**
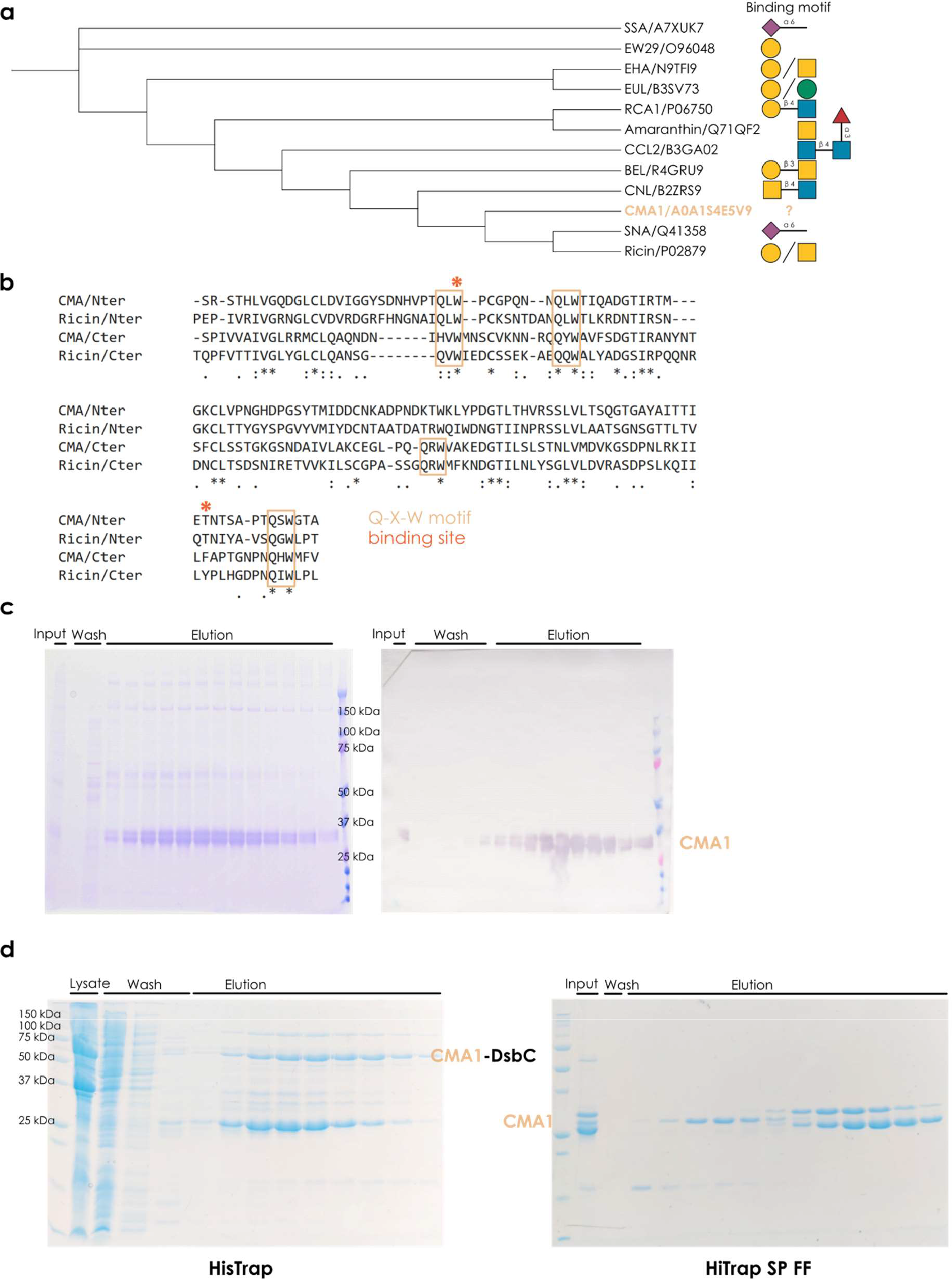
Characterizing a new lectin from the melon *Cucumis melo*. **a)** Evolutionary relationships of common R-type lectins. For a range of representative R-type lectins, we aligned their protein sequences via MUSCLE^21^ and built a neighbor-joining tree with the resulting alignment distances, which is shown as a cladogram. For each protein, we only used the lectin domain, as annotated by UniProt or InterPro. For each protein, a representative binding specificity, based on literature reports, is provided. **b)** Similarity of the two CBM13 domains in CMA1. Using MUSCLE to align the N-terminal (34-158) and C-terminal domains (162-286) of CMA1 and ricin (321-448 and 451-575), we indicated the position of the conserved Q-x-W motif in R-type lectins. **c)** Recombinant expression of CMA1 in mammalian cells. SDS-PAGE and anti-His-tag Western blot of fractions from the expression of CMA1 protein in CHO-S cells. **d)** Recombinant expression of CMA1 in bacteria. SDS-PAGE gels of the His-tag affinity chromatography and cation exchange chromatography from the expression of CMA1 protein in *E. coli* BL21* cells.

We next aligned the individual units of the tandem repeat CBM13 domains, indicated by the N-terminal (34-158) and C-terminal units (162-286) and compared those to the domains of ricin (Fig. 1b). R-type lectins have a characteristic Q-x-W structural motif close to their binding site that is highly conserved^20^. We report that CMA1 largely follows this trend, with three such binding sites in both N- and C-terminal domain, albeit with imperfect overlap. Based on the location of the known binding pocket of the R-type lectin ricin and the respective sequence conservation in CMA1, we postulate binding sites around W^63^ for the N-terminal domain and F^273^ for the C-terminal domain of CMA1.

As binding specificities of melon lectins in general (beyond chitooligosaccharides), and CMA1 in particular, are still unknown, we set out to measure, quantify, and understand the glycan-binding properties of CMA1 in depth, as an archetypal example of melon lectins. For this, we needed to express the lectin recombinantly. As it is a secreted plant protein, we elected to express it in mammalian cell lines, to maximize the chances of a functional protein, due to post-translational modifications that would be lacking in bacteria as well as the oxidative environment of the secretory pathway, as CMA1 exhibits predicted disulfide bridges. A single step of His-tag affinity chromatography was sufficient to yield protein of adequate purity and good yield (∼15 mg of eluted protein from 800 mL of cell culture, Fig. 1c).

In parallel, we also expressed CMA1 in a bacterial expression system, which allowed us to ascertain whether binding was influenced by lectin glycosylation. The full-length mature protein (6-264) and individual N- or C-terminal domains were expressed using a N-terminal fusion comprising DsbC and a hexa-His tag, cleavable by TEV (*Tobacco etch virus*) protease. Despite the presence of the DsbC signal peptide, we did not observe periplasmic localization, and all proteins were instead purified from the cytoplasm. Ni-NTA affinity chromatography followed by TEV protease cleavage of the fusion construct and subsequent reverse Ni-NTA affinity chromatography resulted in significant co-purification of *E. coli* contaminants, necessitating an extra purification step, where cation exchange chromatography allowed us to obtain pure fractions of CMA1^6-291^. Of note, this additional purification step was not necessary for the purification of the CMA1 N-terminal domain. (Fig. 1d). Expression of the CMA1 C-terminal domain did not yield pure and homodisperse enough protein for further analyses.

### *Cucumis melo* agglutinin binds C2-substituted, beta-linked galactose

We then set out to answer the question whether CMA1 was a functional lectin and, if yes, what its binding specificity was. The standard approach to elucidate lectin binding specificity is via glycan array experiments. Here, soluble lectin is added to, traditionally, immobilized glycans and bound lectin is quantified via fluorescence scanners, which can be paired with glycan information due to the known arrangements of immobilized glycans on the plate. To cover the broadest possible sequence space, we tested our eukaryotically produced CMA1 protein against the two largest glycan arrays, at the National Center for Functional Glycomics (NCFG, Fig. 2a) and the Glycosciences Laboratory at Imperial College London (ICL, Fig. 2b). We note that, together, this encompasses 1,046 unique glycans, spanning all major glycan classes and substantial taxonomic diversity. Next to these unique sequences, even more effects stem from a variety of linkers with which these molecules are immobilized.

**Figure 2.**
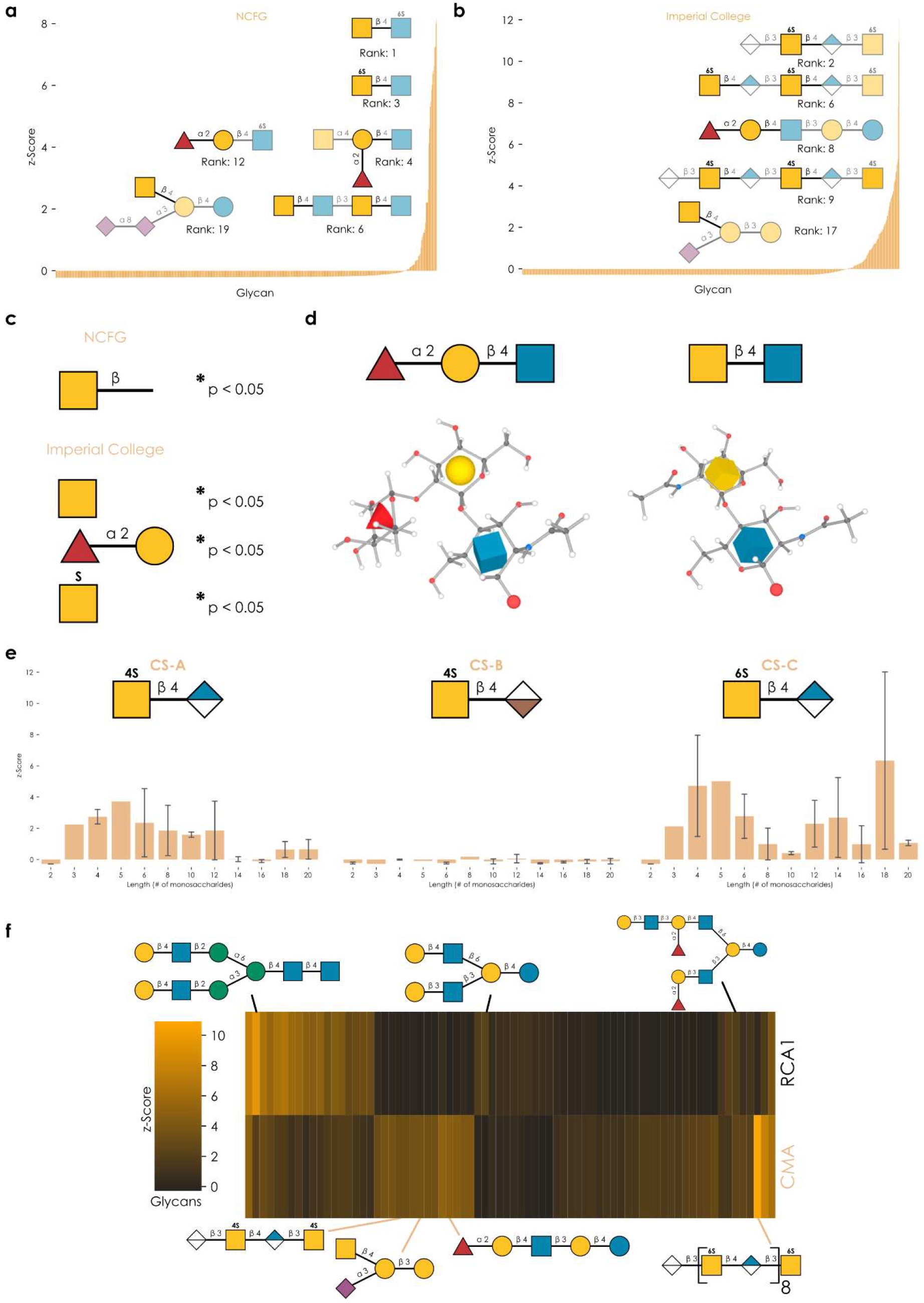
Characterizing the binding specificity of CMA1. **a-b)** Lectin produced in mammalian cells was analyzed on the NCFG array (a) and the ICL array (b). Representative structures bound by CMA1 are shown via the Symbol Nomenclature For Glycans (SNFG), drawn with GlycoDraw^22^. Everything except the purported binding motif is shown with added transparency. **c)** Enrichment analysis of glycan array data. For both NCFG and ICL array data, we used the *get_pvals_motif* function from glycowork^23^ (version 0.8.1) with the keywords ‘terminal’ and ‘exhaustive’, to obtain significantly enriched motifs. *p < 0.05. **d)** Common binding motif on the atomic level. Glycan 3D structures for the binding motifs were obtained from the GLYCAM web server^24,25^. **e)** Binding of CMA1 to glycosaminoglycans. We grouped chondroitin sulfate (CS) types (A, B, and C) and plotted CMA1 binding against CS chain length. Shown are mean values with their 95% confidence interval. **f)** Comparing CMA1 and RCA1 binding. Glycans with a z-score of at least 0.5 in at least one lectin were retained and plotted as a hierarchically clustered heatmap via the *get_heatmap* function of glycowork. Representative glycans are shown.

In general, we observed two binding preferences that were strongly enriched among bound sequences: glycans containing Fucα1-2Gal epitopes and glycans containing terminal GalNAc residues (Fig. 2c). Amongst the bound sequences, these substructures occurred in many different contexts, such as blood group H, LacdiNAc, or the Sd^a^ motif, and particularly in sequences resembling *O*-glycans, milk oligosaccharides and glycosphingolipids. At first glance, these two binding specificities may seem unconnected, indicating a rather broadly binding lectin. However, we noticed that the commonality of these two epitopes is hidden in the IUPAC-condensed nomenclature: Both substructures exhibited a bulky substituent on C2 of galactose, either a fucosyl (Fucα1-2Gal) or *N*-acetyl (GalNAc) moiety (Fig. 2d). We thus conclude that CMA1 is highly specific for C2-substituted galactose. We further argue for a preference for a beta-linked epitope as, while we do observe binding to structures containing α-linked GalNAc, the binding to their β-linked counterparts was generally stronger (e.g., GalNAcα: 1.57 vs. GalNAcβ: 2.21, in z-scores). In part, this is reminiscent to the LacdiNAc binding specificity of *Clitocybe nebularis* lectin (CNL; Fig. 1a)^26^.

An important finding from the ICL array was that CMA1 exhibited robust binding to glycosaminoglycans (GAGs; Fig. 2e), in particular chondroitin sulfate (CS) C and A. Given the preference for terminal binding epitopes described above, the question naturally arose how the binding to these longer-chain glycans functions. On the ICL array, CS sequences are typically capped with hexuronic acid derivatives on their non-reducing end and thus do not provide terminal GalNAc epitopes for binding. Further, while CMA1 did also bind to GalNAc-terminated GAGs (e.g., CSC-5, CSA-5), we measured higher binding to similar GAGs without the terminal GalNAc in several cases (Fig. 2d-e). We thus posit a binding to internal GalNAc epitopes for the case of GAG binding, potentially mediated by several binding sites.

This argument is strengthened by the observation that the highest observed binding to CSC and CSA was not with the shortest sequences and required at least three repeats, with longer sequences such as CSC-18 even exhibiting the highest binding on the entire array (although we note that the longest GAG sequences were not generally the best binders, potentially hinting at steric clashes or density effects). Another supporting finding can be seen in the fact that CSB (exhibiting iduronic acid in α-configuration, rather than its epimer, glucuronic acid, in β-configuration) did show virtually no binding to CMA1, further arguing for contacts of the GAG chain with the binding site. Lastly, we note that both CSC and CSA contain sulfated GalNAc, which, together with the observation of GalNAc6Sβ1-4GlcNAc as one of the highest binders on the NCFG array, leads us to speculate that sulfation further enhances CMA1 binding, a pattern that has been observed for several lectins^27^.

Overall, this characterized binding specificity seemed distinct from other R-type lectins and we thus further compared it to a typical R-type lectin, *Ricinus communis* agglutinin (RCA1), on the ICL array. Canonically, RCA1 binds β-linked terminal galactose residues, which is generally what we also found in our array experiments, with Galβ in various substructures and glycan types, particularly in those with multiple branches (Fig. 2f). At best, the same sequences showed weak binding to CMA1, as they lacked a C2-substitution (Fig. S1). Conversely, CMA1-favored sequences, containing Fucα1-2Gal or GalNAc epitopes, were on average not bound by RCA1 (the exception being sequences in which there was an additional free Galβ terminus). Similarly, most chondroitin sulfate probes were not bound by RCA1. This gives rise to the conclusion that CMA1 does not merely tolerate but rather actively and strongly prefer C2-substitution, while RCA1 does not even tolerate these substitutions. Interestingly, we also find that fucosylation of the GlcNAc residue (as in Lewis antigen motifs) completely abrogates CMA binding (Fig. S1), despite the presence of Fucα1-2Gal, likely due to steric clashes in the binding pocket. We thus conclude that the binding profile of CMA1 is distinct from that of the typical R-type lectin RCA1 and unusual for a R-type lectin in general. We also note that the flexibility of accommodated C2 substituents (from *N*-acetyl moieties to whole monosaccharides), could make CMA1 an interesting candidate for probing synthetically produced glycans with novel substituents.

It is of course interesting to speculate about the physiological role of CMA1 in melons, yet hard to probe. It is noteworthy, however, that the glycan types in which its preferred binding motifs occur (*O*-glycans, milk glycans, GAGs) are absent from most plants, including melons. We thus hypothesize that the role of this lectin might be to recognize non-self epitopes, such as for protection against pathogens, which is a common function in plant lectins^3^.

### Validating binding in solution and assessing binding affinity

As CMA1 both exhibited multiple binding sites and robust binding to blood group epitopes (H-antigen), we hypothesized that it would be capable of agglutinating red blood cells, justifying its new name. Testing protein recombinantly produced in mammalian cells, incubation with rabbit erythrocytes indeed resulted in moderate agglutination (Fig. 3a), which also demonstrated the binding to these glycan substructures in a physiological context.

**Figure 3.**
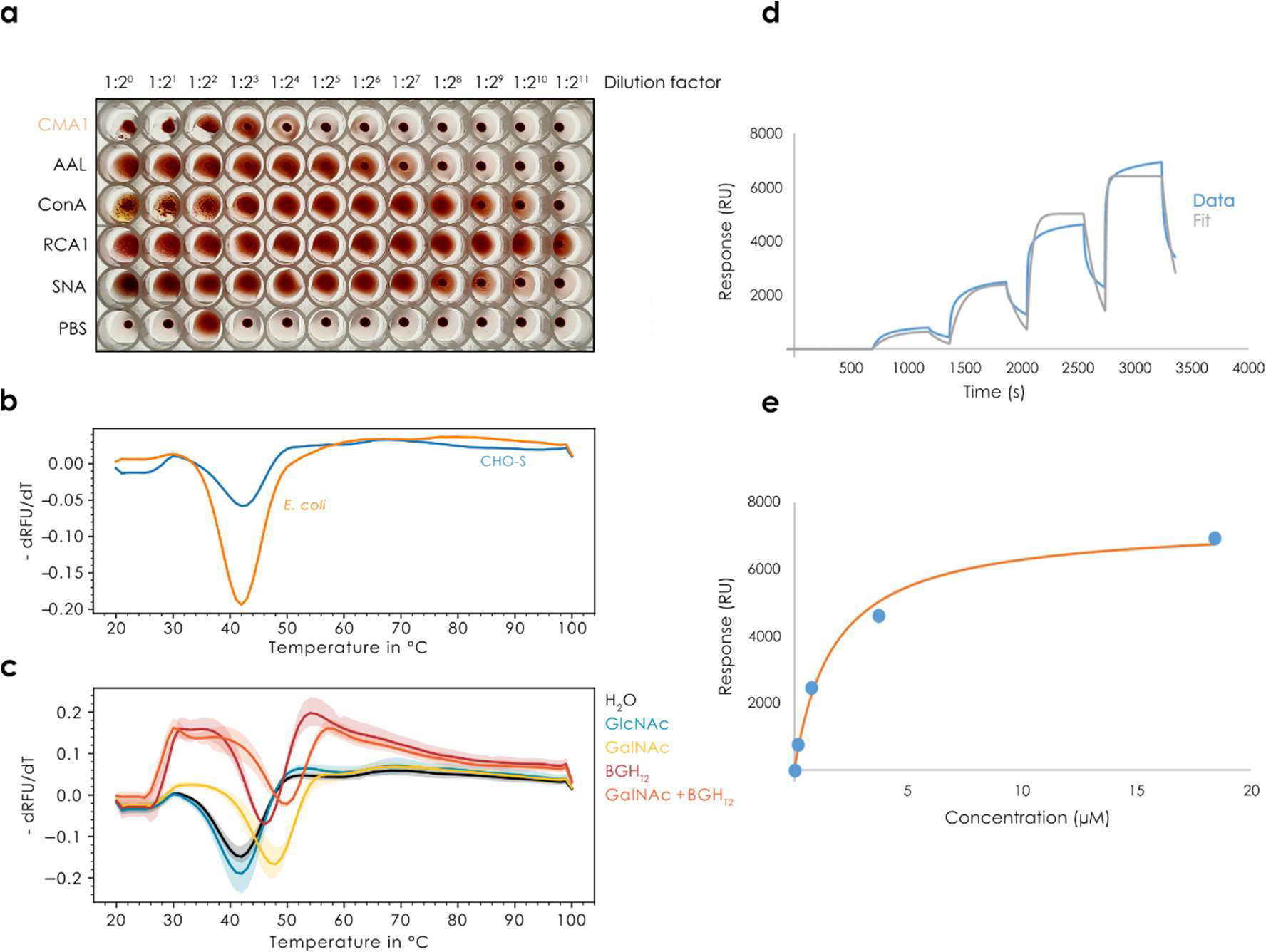
Assessing and quantifying in-solution binding of CMA1. **a)** Erythrocyte agglutination assay. Using rabbit red blood cells, CMA1 protein recombinantly produced in mammalian cells was used in a two-fold dilution series to measure its ability to agglutinate erythrocytes, compared to other lectins, such as AAL, ConA, RCA1, and SNA-I, as well as a PBS negative control. **b-c)** Thermal shift assay. After comparing the melting curves of CMA1 produced in mammalian cells (CHO-S) and bacteria (*E. coli*), we incubated the bacterially produced CMA1 with GlcNAc, GalNAc, and H type 2 blood group antigen (BGHT2; Fucα1-2Galβ1-4GlcNAcβ1-3Gal) and measured a denaturation curve to assess shifts in melting temperature, n = 3 (c). **d-e)** SPR analysis of CMA1 binding to a GalNAc chip with single-cycle kinetics and affinity measurement at the equilibrium, n = 2.

To further strengthen the case for CMA1 binding glycans in solution, and corroborate its binding specificity with orthogonal methods, we used a thermal shift assay. Herein, the binding of ligands is assessed by a stabilization of the protein, measured by a denaturation curve. Both the protein produced in mammalian and in bacterial cells exhibited similar melting temperatures here, of approximately 42°C (Fig. 3b). Then, we tested the binding of CMA1 to GlcNAc, GalNAc, and H type 2 blood group antigen (BGH_T2_; Fucα1-2Galβ1-4GlcNAcβ1-3Gal; Fig. 3c). This resulted in clear melting points shifts for both GalNAc and BGH_T2_ to up to 50°C, yet importantly not for GlcNAc, demonstrating both binding in solution and a further confirmation of the binding specificity obtained by the array experiments.

Next, we set out to quantify the binding affinity of CMA1 to its ligands. Lectins often only exhibit weak to moderate binding affinities, which is somewhat ameliorated by an increased avidity on the side of the lectin but also a dense presentation of the bound glycan epitope on the cell surface. We therefore used surface plasmon resonance (SPR) to derive binding constants for the interaction between CMA1 and GalNAc. A single cycle kinetics approach was applied, resulting in a measured K_D_ of 1.66 +/- 0.08 μM (Fig. 3d-e). Inhibiting binding of CMA1 to the GalNAc chips through a dilution series of lactosamine via multicycle kinetics allowed us to derive an IC_50_ of 1.4 μM (Fig. S2a-b). No inhibition was observed with chondroitin 6-sulfate tetrasaccharide and only very weak inhibition for BGH_T2_ but no IC_50_ could be determined as we could not increase the concentration to reach the plateau. For the recombinant CMA1-Nter, no binding could be observed on the GalNAc chip. This suggests either avidity effects in conjunction with the C-terminal domain or a high-affinity site on the C-terminal domain, giving rise to the measured K_D_ of the full-length protein. Still, we were able to measure the affinity of CMA1-Nter to GalNAc in solution by isothermal calorimetry (ITC), obtaining a K_D_ of 940 μM, confirming the low affinity (Fig. S2c-d).

### Structural insights from the N-terminal domain of CMA1

Given the unusual binding specificity exhibited by CMA1, we were intrigued to elucidate the molecular mechanism that would enable the specific binding of C2-substituted galactose. The natural hypothesis here would be the creation of an additional pocket in the 3D structure of the binding site, accommodating the additional substituent at C2. However, as we observed little to no binding to unsubstituted galactose, we rather hypothesized the existence of specific interactions made with the C2-substituents, that did not exist in other R-type lectins such as RCA1. To determine this, we set out to resolve the detailed three-dimensional structure of CMA1 via X-ray crystallography.

We obtained several hits for the full-length protein after sparse screening using a crystallization robot at the HTX platform, EMBL, Grenoble. Pill-shaped crystals obtained in conditions containing a high salt concentration, in particular ammonium sulphate (Fig. S3), did not give rise to any diffraction. Multiple layer plate or needles clusters were obtained in the presence of PEGs, but only showed weak diffraction (∼3.5Å). Finally, in the presence of 20% PEG 8K, 0.2 M MgCl_2_, and 0.1 M Tris HCl pH 8.5, single diamond-shaped crystals were obtained in 1-2 days for the N-terminal domain (Fig. S3). The crystals diffracted to near-atomic resolution and the CMA1-Nter structure was solved in complex with lactosamine at 1.3 Å and GalNAc at 1.55 Å resolution (see data and refinement statistics in Table 1). All residues of the N-terminal construct (Val^6^ to Asp^132^) could be modelled, and unambiguous electron density permitted us to locate and model four cation binding sites (three in each structure) and one sugar binding site (Fig. 4a-b, Fig. S4).

**Table 1.** Data collection and refinement statistics.

| **Complex** | **CMA1-Nter-LacNAc** | **CMA1-Nter-GalNAc** |
| --- | --- | --- |
| **Data collection** |  |  |
| Beamline | Soleil PX1 | Soleil PX2 |
| Wavelength (Å) | 0.97856 | 0.98011 |
| Space group | *I*2 | *I*2 |
| Cell parameters – *a*, *b*, *c* (Å) | 36.70 36.78 94.79 | 36.61 36.86 94.81 |
| –α, $\beta$, $\gamma$ (°) | 90.00 99.24 90.00 | 90.00 99.17 90.00 |
| Protein chains in a.u. | 1 | 1 |
| Resolution (Å)<sup>a</sup> | 46.78- 1.32 (1.34-1.32) | 35.68-1.55 (15.8-1.55) |
| CC1/2 (%)<sup>a</sup> | 99.9 (96.9) | 99.8 (85.7) |
| *R*<sub>merge</sub> (within I+/I-) <sup>a</sup> | 0.055 (0.369) | 0.052 (0.496) |
| *R*<sub>meas</sub> (within I+/I-) <sup>a</sup> | 0.059 (0.400) | 0.064 (0.618) |
| *R*<sub>pim</sub> (within I+/I-) <sup>a</sup> | 0.022 (0.153) | 0.037 (0.364) |
| Mean I/$\sigma$ (I) <sup>a</sup> | 25.2 (5.7) | 14.4 (2.9) |
| Completeness (%)<sup>a</sup> | 99.8 (96.0) | 99.7 (99.9) |
| Number reflections <sup>a</sup> | 399970 (18410) | 95115 (4581) |
| Number of unique reflections <sup>a</sup> | 29695 (1434) | 18279 (911) |
| Multiplicity<sup>a</sup> | 13.5 (12.8) | 5.2 (5.0) |
| Wilson *B*-factor (Å<sup>2</sup>) | 14.1 | 19 |
| **Refinement** |  |  |
| Resolution (Å) | 46.78 -1.32 | 35.69-1.55 |
| No. reflections / No. free reflections | 28192 / 1503 | 17373 / 905 |
| *R*<sub>work</sub>/*R*<sub>free</sub> (%) | 14.35 / 18.58 | 16.3 / 20.4 |
| R.m.s. Bond lengths (Å) | 0.0130 | 0.0127 |
| Rmsd Bond angles (°) | 1.721 | 1.893 |
| Rmsd Chiral (Å<sup>3</sup>) | 0.097 | 0.092 |
| No. atoms / Bfac (Å<sup>2</sup>) |  |  |
| Protein | 1029 / 15.1 | 985 / 19.95 |
| Ligand | 26 / 20.3 | 30 / 22.3 |
| Cadmium | 3 / 21.9 | 3 / 27.0 |
| Waters | 248 / 28.7 | 176 / 31.8 |
| Ramachandran Allowed (%) | 100 | 100 |
| Favored (%) | 99 | 100 |
| Outliers | 0 | 0 |

<sup>a</sup> Values in parenthesis refer to the highest-resolution shell

**Figure 4.**
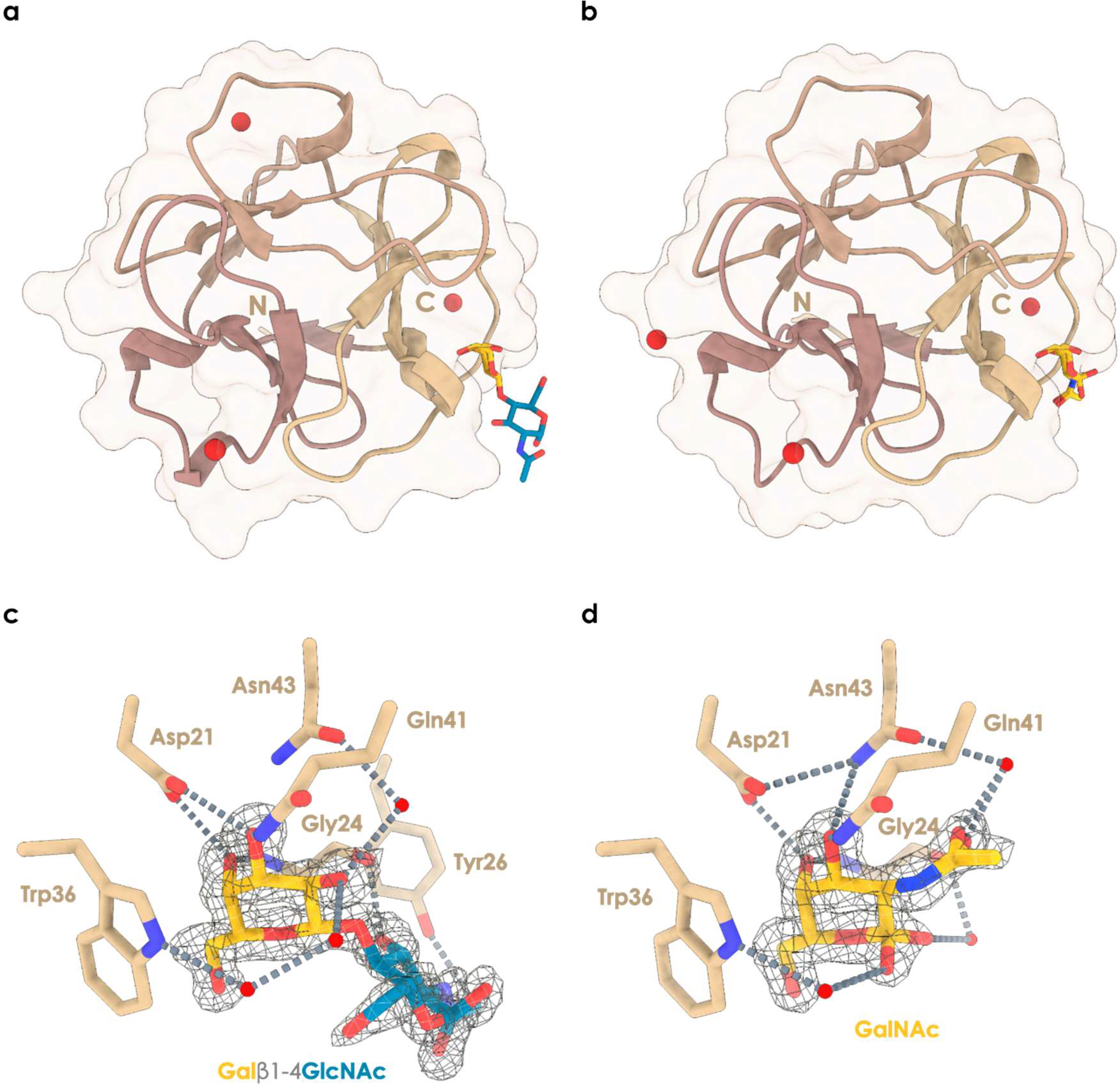
Structural insights into the binding mechanism of CMA1. **a-b)** Overall representation of the N-terminal domain of CMA1 in complex with LacNAc (Galβ1-4GlcNAc) (a, PDB ID 8R8A) or GalNAc (b, PDB ID 8R8C). Trefoil repeats are colored differently, and cadmium ions are represented as red spheres. **c-d)** Close-up on the interactions between CMA1 and LacNAc (c) or GalNAc (d), with electron density displayed around the sugar ligands at 1 sigma (LacNAc: 0.47 eÅ^-3^, GalNAc: 0.415 eÅ^-3^). Water molecules are indicated by red spheres and interactions by proximal residues are indicated by broken lines.

The complexed structures allowed us to shed light on the arrangement of the ligand in the binding site (Fig. 4c-d). While lectins such as CMA1 typically can present three binding pockets in their CBM13 domain, we hypothesized that the N-terminal half of CMA1 would in fact only exhibit two functional binding sites. However, only the alpha site was found occupied with a carbohydrate here. It is found in a shallow groove, supporting our data on the lack of a distinct distal binding specificity. We report a tight coordination of the O3 and O4 hydroxyls of the galactose residue involving Asp^21^, Asn^43^, and Gln^41^ side chains, as well as the Gly^24^ main chain nitrogen. Stacking and hydrophobic interactions are made between the aromatic ring of Trp^36^ and the alpha face of the ring as well as the hydroxymethyl moiety of the galactose residue, additionally ensuring specificity for galactoside over glucoside as an equatorial conformation of the O4 hydroxyl would lead to steric clashes and loss of strong hydrogen bonding.

In structure 8R8A, the GlcNAc residue did not seem to engage in extensive interactions, with only a hydrogen bond between the *N*-acetyl moiety and the main chain oxygen of Gly^24^ and hydrophobic interaction with the aromatic ring of Tyr^26^ (Fig. 4c). Further, beyond the C2 position of galactose, a cavity filled with coordinated water molecules hinted at the binding mode for C2-substituted galactose. Notably, the seemingly inactive beta site was found to be occupied by a cadmium ion (Fig. S4), supporting our ITC and SPR data where no multivalent binding effects were observed for the single-domain N-terminal construct.

In structure 8R8C, the *N*-acetyl group of GalNAc extended beyond C2 into the cavity noted in 8R8A. While no direct interactions with the protein backbone were observed, we found one water molecule to mediate hydrogen bonding between the oxygen of the *N*-acetyl group and the Asn^43^ side chain oxygen (Fig. 4d). Both GalNAc anomers could be observed, showing interaction through water molecule coordination with the Trp^36^ ring nitrogen or the Gly^24^ main chain oxygen.

## Conclusion

Our work presents a substantial exploration of the binding specificity and mechanism of the hitherto uncharacterized lectin CMA1 from melons. The binding specificity of CMA1, C2-substituted galactose that is preferentially presented in a β-configuration, enables it to bind to a range of biologically relevant epitopes, such as LacdiNAc, Sd^a^, blood group H, and chondroitin sulfate motifs. Further, the inhibition of binding by the presence of Lewis antigen motifs additionally narrows it binding specificity. Our binding data and structural information lead us to the conclusion that crucially positioned asparagine residues facilitate this unusual binding specificity that delineates CMA1 from typical R-type lectins such as RCA1. Together, these results advance our knowledge of R-type lectins in general, and the range of their binding specificities, but also our knowledge of melon lectins in particular, which has remained limited so far. Further experiments are still required to determine the role of the C-terminal domain, as well as the physiological function of the full-length CMA1 protein.

## Experimental

### Recombinant protein expression

For mammalian expression, the gene for CMA1 (A0A1S4E5V9) was synthesized with human-optimized codons and a C-terminal hexa-His tag (GSHHHHHH). We then cloned this gene into a pCI backbone (U47119; Promega GmbH) for expression in mammalian cells under a constitutive CMV promoter. Then, the Mammalian Protein Expression core facility at the University of Gothenburg transfected this plasmid into FreeStyle™ CHO-S Cells (Cat nr R80007, ThermoFisher Scientific). Cells were cultured in Freestyle™ CHO medium at 37°C in 5% CO_2_ in Optimum Growth™ flasks (Thomson instrument company) at 130 rpm in a Multitron 4 incubator (Infors) and transfected at 2x10^6^ cells/mL using FectoPro transfection reagent (Polyplus). Protein-containing culture supernatant (0.8 L) was harvested after 120 hours, filtered using Polydisc AS 0.45 μm (Whatman, Cytiva) and loaded onto a 5 mL HisExcel column (GE healthcare) at 5 mL/min. The column was washed with 10 mM phosphate-buffered saline (Medicago), 500 mM NaCl and 50 mM imidazole before elution of the protein using the same buffer with a gradient from 50 mM to 500 mM Imidazole (G-Biosciences) over 15 column volumes. Pooled fractions were concentrated using Vivaspin concentrators (MWCO 10 kDa, Sartorius Stedim), passed over a HiPrep 26/10 desalting column (GE Healthcare) in phosphate-buffered saline (Medicago) and finally concentrated again.

For bacterial expression, the gene of CMA1 (33-291, corresponding to residues 6-291 of the mature protein) with optimized codons for *Escherichia coli* was synthesized flanked by *Nco*I and *Xho*I restriction sites where L^6^ was mutated to valine. The gene was inserted in the homemade plasmid pET40b-TEV where the enterokinase cleaving site was replaced by a TEV cleavage site by site directed mutagenesis. This plasmid was obtained by PCR using pET-40b(+) (Novagen, Merck, #70091) as template and the following primers: forward (<u>GCCCAGATCTGGGTACC</u>GAAAACCTGTATTTTCAGGGCGCCATGGCGATATCGG) and reverse (<u>GGTACCCAGATCTGGGC</u>TGTCCATGTGCTGGC) with complementary sequence underlined. The PCR was performed using PrimeSTAR DNA polymerase (Takara #TAKR045A) then the product was digested by *Dpn*I, and finally transformed in NEB5α strain (New England Biolabs, #C2992H). Both gene and vector were digested by *Nco*I and *Xho*I restriction enzymes (New England Biolabs) prior to purification on agarose gel using Monarch Gel extraction kit and supplier instructions (New England Biolabs, #T1020S) and ligation using the DNA ligation kit, Mighty Mix (Takara, #6023), at room temperature to form the pET40b-TEV-CMA11 plasmid.

The N-terminal domain of CMA1 (6-132 in mature protein) was amplified by PCR using the following primers (Forward: ACG<u>CCATGG</u>TGAGCCGTTCTACGC and Reverse: ATAT<u>CTCGAG</u>TTAATCTG CCGTACCCCAGGATTGTGTAGG) and pET40b-TEV-CMA1 plasmid as template. Similarly, the C-terminal domain of CMA1 (136-264 in mature protein) was amplified by PCR using the subsequent primers (Forward: ATT<u>CCATGG</u>GTCCGATTGTGGTTGCCATTGTTGG and Reverse: ACAC<u>CTCGAG</u>TTAGGGTTTGTACTGTGTCACGAACATCC. The primers contained the restriction sites (underlined) *Nco*I (sense) and *Xho*I (antisense) on their 5′-ends for further sub cloning. PCR was performed using PrimeSTAR DNA polymerase. The purified PCR fragment of 395 bp was digested by *Nco*I and *Xho*I restriction enzymes, then ligated into pET40b-TEV vector, and finally transformed in NEB5α strain to form the pET40b-TEV-CMA1-Nter and pET40b-TEV-CMA1-Cter plasmids. All plasmids and new vectors were verified by sequencing (Eurofins Genomics, Ebersberg, Germany). Primers were purchased from Eurofins Genomics (Ebersberg, Germany).

*E. coli* BL21*(DE3) [Invitrogen, #C601003] cells were transformed by heat shock at 42°C with pET40b-TEV-CMA1 and Tuner(DE3) [Novagen, #70623] cells with pET40b-TEV-CMA1Nter prior pre-culturing in lysogeny broth (LB) [Invitrogen, #12780052] media containing 25 μg/mL kanamycin [Euromedex, #UK0015-A] at 37°C, 180 rpm overnight. Then, 1 L LB medium supplemented with 25 μg/mL kanamycin was inoculated with 25 mL of the pre-culture and incubated at 37°C, 180 rpm. When OD_600nm_ reached 0.4, the temperature was lowered to 16°C and when OD_600nm_ reached 0.8, protein expression was induced by the addition of 0.1 mM isopropyl β-d-thiogalactoside (IPTG) [Euromedex, #EU0008-C]. After 20 hours, the cells were harvested by centrifugation at 5,000 x *g* for 10 min at 4°C.

For purification of bacterial recombinant CMA1, each gram of cell pellet was resuspended with 5 mL of buffer A (20 mM Tris-HCl pH 7.5, 500 mM NaCl). After addition of 1 μL of Denarase (C-LEcta GmbH, #20804) and moderate agitation on a rotating wheel during 30 min at room temperature, cells were lysed using a cell disruptor (Constant Systems Ltd, UK) under a pressure of 2.5 kbar. The lysate was cleared by centrifugation at 24,000 x *g* for 30 min at 4°C and passed through a 0.45 μm syringe filter prior to affinity chromatography purification using 1 mL HisTrap™ HP column (Cytiva) preequilibrated with buffer A and an NGC chromatography system (Bio-Rad). After loading the cleared lysate, the column was washed with Buffer A + 50 mM imidazole (Sigma-Aldrich, Merck, #56749) to remove all contaminants and unbound proteins. CMA1 was eluted by a 20 mL linear gradient from 50 mM to 500 mM imidazole in Buffer A. The fractions were analyzed by SDS-PAGE with 15% gel and those containing CMA1 were collected and deprived of imidazole by buffer exchange in buffer A using a Macro and Microsep Advance Spin 3 kDa MWCO centrifugal filter (Pall). The N-terminal His-tag was removed by TEV cleavage in the presence of 1 mM EDTA (Euromedex, #EU0084.B) overnight at 10°C, using a TEV:CMA1 ratio of 1:50. TEV was prepared in-house. The protein mixture was then purified on a 1 mL HisTrap column, where pure CMA1 protein was collected in the flowthrough and column wash. Full-length CMA1 (6-291) was purified from remaining *E. coli* contaminants using a 1 mL HiTrap™ SP Sepharose FF column (Cytiva) preequilibrated with 50 mM sodium acetate pH 5.5. After loading, the column was washed, and CMA1 eluted by a 20 mL linear gradient from 0 mM to 700 mM NaCl in 50 mM sodium acetate pH 5.5. The protein was concentrated and the buffer exchange to 20 mM HEPES pH 8, 100 mM NaCl using a 3 kDa MWCO centrifugal filter and stored at 4°C.

For CMA1-Nter, the same protocol was followed, with the following changes: Purification was made by gravity using 1 mL of Ni Sepharose High Performance resin (Cytiva, #17.5268.01) and an Econo-Pac® Chromatography Column (Bio-Rad, #7321010). Buffer A was exchanged to buffer B (20 mM Tris-HCl pH 8.0, 500 mM NaCl, 500 mM urea, and 5 mM imidazole). Washing steps were performed using buffer B and buffer B containing 50 mM imidazole. Elution was performed using buffer B plus 250 mM imidazole. The buffer was exchanged to 20 mM HEPES pH 7.5, 100 mM NaCl by three times 10x dilution and the sample was concentrated to at least 1 mg/mL using a 3 kDa MWCO centrifugal filter prior to TEV cleavage.

### Glycan array experiment

CFG array. For the CFG array, data was collected by the National Center for Functional Glycomics (NCFG) at Beth Israel Deaconess Medical Center, Harvard Medical School. For experiments, a standard binding buffer (20 mM Tris-HCl pH 7.4, 150 mM NaCl, 2 mM CaCl_2_, 2 mM MgCl_2_, 0.05% Tween 20, 1% BSA) was used. CMA1 binding was probed by incubation with a penta-His-488 antibody (5 μg/mL). CMA1 was tested in two concentrations (5 and 50 μg/mL) on Version 5.4 of the printed CFG array, consisting of 585 printed glycans in replicates of six. Results from replicates were combined as average RFU (raw fluorescence unit). For this average, the highest and lowest value was removed for each glycan, mitigating the effects of outliers. The results can be found in Supplementary Table 1.

Imperial College array. For experiments, a standard binding buffer (10 mM HEPES, 150 mM NaCl, 1% BSA, 0.02% casein blocker (Pierce), 5 mM CaCl_2_) was used. CMA1 was tested at 100 μg/mL for one hour on the large screening panel of the Glycoscience Laboratory at Imperial College London, consisting of 866 lipid-linked glycans. Then the detecting solution composed of anti-polyHistidine (Sigma-Aldrich, Merck, SAB4200620) and biotin anti-mouse IgG (Sigma-Aldrich, Merck, B7264) antibodies (10 μg/mL, precomplexed in a ratio of 1:1) was overlaid onto the arrays for one hour. The final detection was with a 30-minute overlay of streptavidin-Alexa Fluor 647 (Molecular Probes) at 1 μg/mL. The microarray slides were scanned with GenePix 4300A scanner instrument (50% laser power at PMT 350) and the image analysis (quantitation) was performed with GenePix^®^ Pro 7 software. The results can be found in Supplementary Tables 2-3.

### Agglutination assay

The hemagglutinating activity of CMA1 was determined in V-bottom 96-well plates by a 2-fold serial dilution procedure in PBS using rabbit red blood cells (Atlantis France). 25 μL of 4% erythrocyte suspension was added to an equal volume of the sample and the mixture incubated for 60 minutes at room temperature. Starting concentrations were: CMA1 0.6 mg/mL, AAL 0.5 mg/mL, ConA 2.5 mg/mL, RCA1 2.5 mg/mL, SNA 0.5 mg/mL.

### Thermal shift assay

Thermal shift assays (TSA) were performed using a Mini Opticon Real Time PCR machine (BioRad). 0.6 mg/mL protein in PBS was mixed with SYPRO Orange (Sigma-Aldrich, Merck, #S5692) and glycan ligand (10 mM GalNAc; Carbosynth, #MA04390; 10 mM GlcNAc, Carbosynth, #MA00834; 10 mM blood group H type 2 tetrasaccharide; Elicityl, GLY032-2) in a total reaction volume of 25 μL. The temperature was raised by 1°C/min from 25°C to 100°C and fluorescence readings were taken at each step.

### Surface plasmon resonance (SPR)

Experiments were performed using a Biacore X100 instrument (Cytiva) at 25°C in HBS-T running buffer (10 mM HEPES pH 7.4, 150 mM NaCl and 0.05% Tween 20). Biotinylated PAA-GalNAc (Lectinity, GlycoNZ, #0031-BP) was immobilized on CM5 chips (Cytiva # BR100012) that were coated previously with streptavidin (Sigma-Aldrich, Merck, #S4762), following standard protocol. Biotinylated GalNAc was diluted to 2 μg/mL in HBS-T before being injected into one of the flow cells of the chip. Immobilization level of 900 response units (RU) was obtained. A reference surface was always present in flow cell 1, allowing for the subtraction of bulk effects and non-specific interactions with streptavidin. The mammalian-produced CMA1 was injected in single cycle kinetic over the flow cell surface at 10 μl/min at increasing concentrations with a contact time of 500 s. Dissociation was achieved by passing running buffer for two minutes. Surfaces were regenerated with four consecutive 30 s injections of 50 mM NaOH and 1 M NaCl. Binding affinity (K_D_) was measured after subtracting the channel 1 reference (streptavidin only) and subtracting a blank injection (running buffer - zero analyte concentration). Data evaluation and curve fitting was performed using the provided BIACORE X100 evaluation software (version 2.0). Measurements were at least done in duplicate.

Then, to perform competition experiments, nine concentrations of LacNAc (Elicityl, # GLY008) from 10 mM to 0 mM with a dilution coefficient of two supplemented with a fixed concentration of 0.8 μM was injected into the cell surface in multiple cycle kinetic with an association time of 500 seconds and a dissociation time of 12 seconds at a flow rate of 10 μl/min. Surfaces were regenerated with 30 s injections of 50 mM NaOH and 1 M NaCl. IC_50_ was measured using the response at equilibrium for each concentration of competitive sugar that were translated in percentage of inhibition, then plotted against the molar concentration of competitive sugar using the free software “data entry”. The IC_50_ was calculated using https://www.aatbio.com/tools/ic50-calculator.

### X-ray crystallography

All consumables for crystallization and crystal handling were purchased at Molecular Dimensions, Calibre Scientific, Rotherham, UK, unless stated otherwise. CMA1 concentrated at 5.7 or 3.5 mg/mL in 20 mM HEPES pH 8, 100 mM NaCl, and 14 mM GalNAc was subjected to crystallization screening using the robotized HTXlab platform (EMBL, Grenoble, France) with 200 nanoliter sitting drops at 20°C using a 1/1 ratio. Wizard I and II screen (Rigaku) and SaltRX (Hampton Research) screens were used and led to more than 30 hits after one to three days. Pill-like crystals were obtained with high salt concentration that could be reproduced by hand in the laboratory. Plates and needles clusters were obtained with PEG containing solutions. For CMA1-Nter, protein at a concentration of 2.9-3.5 mg/mL was crystallized using hanging drop and vapor diffusion method with a 2 μL drop in 1/1 ratio at 20°C. Bipyramidal single crystals were obtained in one or two days in a solution containing 10-12% PEG Smear Medium, 0.1 M MES pH 6.5, 1X divalent (5 mM of CaCl_2_, MgCl_2_, CsCl_2_, CdCl_2_, NiCl_2_, and Zinc acetate), or 5 mM CdCl_2_, and in the presence or not of 5 mM GalNAc. Cocrystals of CMA1-Nter in complex with lactosamine (Galβ1-4GlcNAc, LacNAc, Elicityl, #GLY008) were obtained by the addition of 5 mM LacNAc to the protein solution and incubation at room temperature for 30 minutes prior to crystallization. Single crystals were mounted in a cryoloop after transfer in a cryoprotectant solution when necessary and flash cooled in liquid nitrogen. For CMA1-Nter-LacNAc complex, a solution of 30% PEG Smear Medium and 5mM CdCl_2_ was used for cryoprotection. Crystal diffraction was evaluated, and data were collected on the Proxima 1 beamline at the synchrotron SOLEIL, Saint Aubin, France using an Eiger 16M detector (Table 1). XDS and XDSME were used to process the data and all further steps were performed using programs of the CCP4 suite version 8.25–27^28–30^. The coordinates of Alphafold^31^ Monomer v2.0 Prediction For Nigrin B-Like (A0A1S4E5V9) were used as a search model to solve the structure of CMA1-Nter by molecular replacement using PHASER^32^. Multiple iterations of anisotropic restrained maximum likelihood refinement using REFMAC 5.8^33^ and manual building using Coot^34^ were performed. Hydrogen atoms were added in their riding positions during refinement and 5% of the observations were set aside for cross-validation analysis. Upon inspection of the electron density maps, carbohydrate moieties were introduced and checked using Privateer^35^. The final model was validated using the wwPDB validation server (https://validate-rcsb-1.wwpdb.org). The coordinates of CMA1 in complex with LacNAc (8R8A) and GalNAc (8R8C) were deposited in the Protein Data Bank (PDB). Structure figures were made using PyMol 2.5.7 and ChimeraX 1.6.

## Funding

This work was funded by a Branco Weiss Fellowship – Society in Science awarded to D.B., by the Knut and Alice Wallenberg Foundation, and the University of Gothenburg, Sweden as well as support from the GLYCONanoPROBES (CA18132) and INNOGLY (CA18103) COST actions awarded to J.L. This work was further supported by the Protein-Glycan Interaction Resource of the CFG and the National Center for Functional Glycomics (NCFG) at Beth Israel Deaconess Medical Center, Harvard Medical School (supporting grant R24 GM137763). This work benefited from access to EMBL HTX lab, which has been supported by iNEXT-Discovery, project number 871037, funded by the Horizon 2020 program of the European Commission. We also acknowledge support from the Mammalian Protein Expression core facility at the University of Gothenburg via the Protein Production Sweden (PPS) framework as well as the synchrotron SOLEIL (Saint Aubin, France) for access to beamline Proxima 1 and 2 (Proposal Number 20210859) and for the technical support of Pierre Legrand and Martin Savko, respectively. The authors would like to thank Iris Lopez and Federico Musso for their technical help in the expression assays of CMA1-Nter and CMA1 purification trials, respectively.

## Notes

### Competing Interest Statement

The authors have declared no competing interest.

