## Supplemental Figures for "Elucidating the Glycan-Binding Specificity and Structure of *Cucumis melo* Agglutinin, a New R-Type Lectin"

**Supplementary Figures**

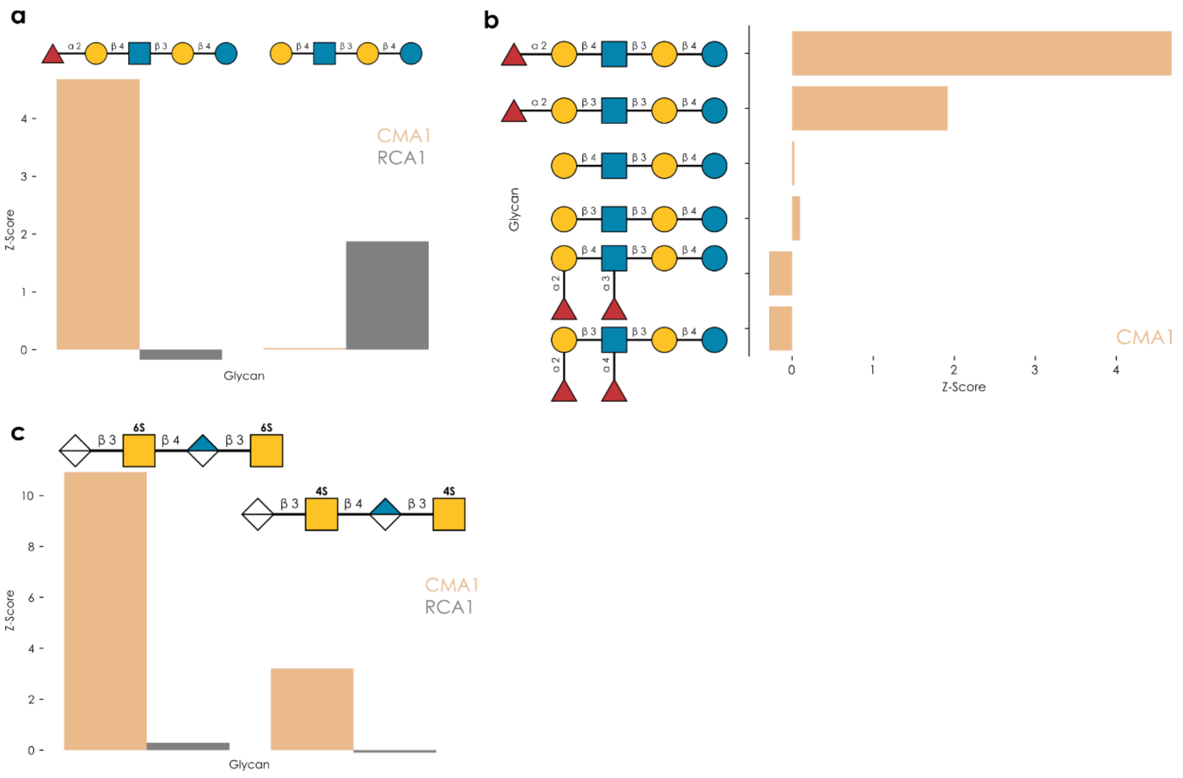


**Figure S1. CMA1 strongly prefers C2-substituted galactose for binding. a-c)** For CMA1 and RCA1, we selected z-scores from the Imperial College London glycan array data to detail the binding specificity of CMA1. This showed that, while RCA1 requires terminal galactose, CMA1 almost requires the extension of the C2 position of galactose (a). Further, extension of the GlcNAc into a Lewis antigen motif completely abrogated binding of CMA1 to its usual binding motif (b). Example glycosaminoglycans (CS-C and CS-A) indicate strong binding of CMA1, but not RCA1, to sulfated GalNAc.

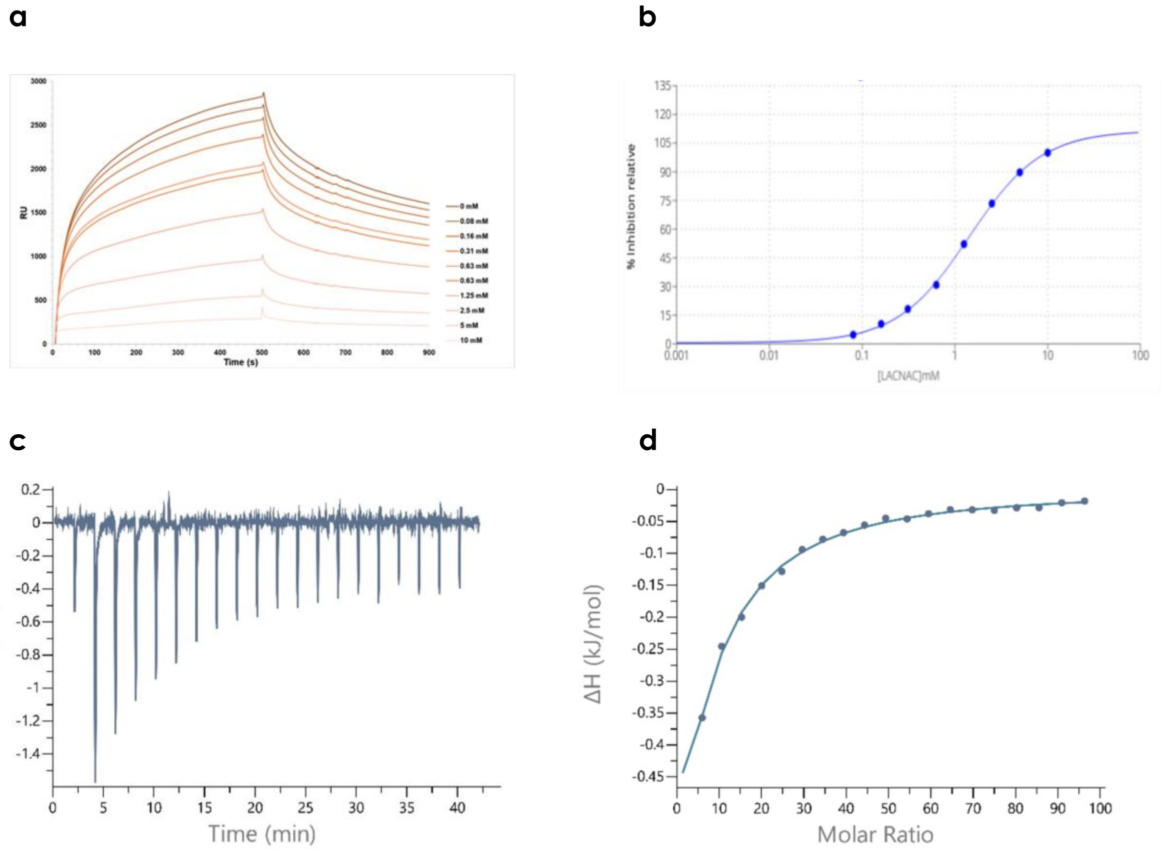


**Figure S2. Assessing and quantifying in-solution binding of CMA1. a-b)** SPR analysis of the inhibition of the binding of CMA1 in the presence of LacNAc: multi-cycle sensograms and IC50 analysis. **c-d)** ITC analysis of the affinity of CMA1-Nter to GalNAc in solution.

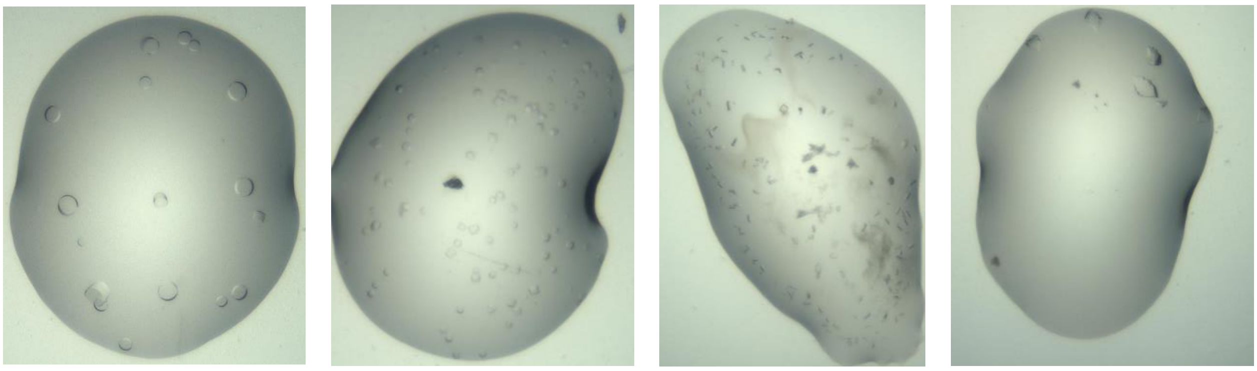

**Figure S3. Crystals of the N-terminal domain of CMA1.** The crystals were obtained, from left to right, with (i) 1.5 M $AmSO_4$, 0.1 M Bis Tris Propane pH 7.0, SaltRX F2; (ii) 4M Ammonium acetate, 0.1 M Tris HCl pH 8.5, SaltRX H10, (iii) 30% PEG 8K, 0.2 M NaCl, 0.1 M Imidazole HCl pH 8, Wizard I-II F12, or (iv) 20% PEG 8K, 0.2 M $MgCl_2$, and 0.1 M Tris HCl pH 8.5, Wizard I+II E3.

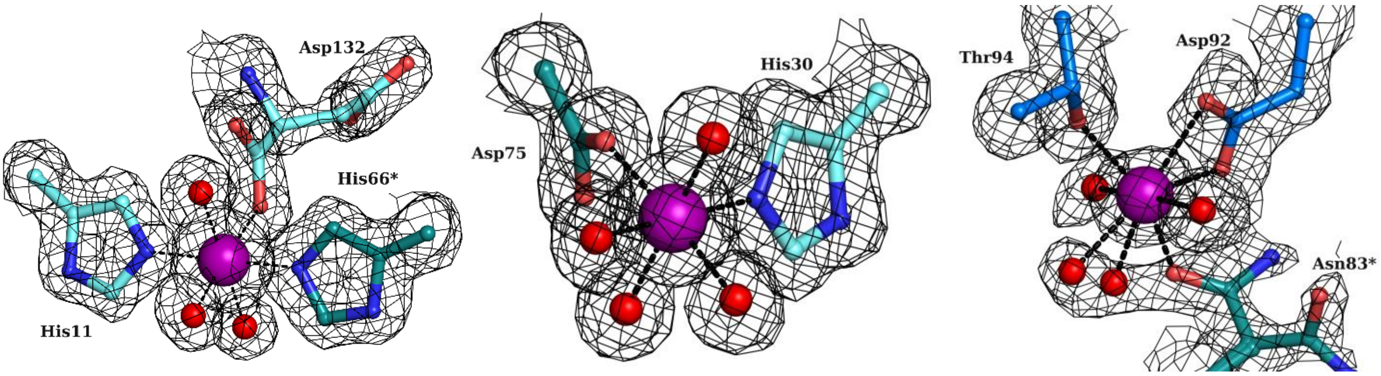


**Figure S4. Metal-binding sites in the N-terminal domain of CMA1.** Electronic density for cadmium ions binding sites of CMA1 represented at 1 sigma (0.47 e Å$^{-3}$). Cadmium is depicted as a purple sphere, waters as red spheres, and amino acids in ball and sticks. Amino acid names with a star represent symmetry related residues.
